# Mitochondria-centered metabolomic map of inclusion body myositis: sex-specific alterations in central carbon metabolism

**DOI:** 10.1101/2024.09.29.615665

**Authors:** Elie Naddaf, Ibrahim Shammas, Surendra Dasari, Xuan-Mai T. Petterson, Eugenia Trushina, Ian R. Lanza

**Author notes:** **Corresponding Author:** Elie Naddaf, MD, Mayo Clinic, Department of Neurology, 200 First Street SW, Rochester, MN 55905.

## Abstract

**Background:** Inclusion body myositis (IBM) is a disease of aging characterized by progressive muscle loss. Despite its positioning at the intersection of aging, mitochondrial dysfunction and chronic inflammation, limited studies have evaluated the underlying metabolic disturbances in IBM.

**Objective:** To investigate the mitochondria-centered metabolomic map of IBM in muscle tissue, highlighting sex-specific differences, and to determine the correlation of the changes in metabolites and gene expression with clinical parameters.

**Methods:** 37 IBM patients and 22 controls without a myopathy were included. All participants had bulk RNA sequencing performed previously. Clinical parameters included age at biopsy, disease duration, manual motor test (MMT) score, and modified Rankin scale (MRS). A complementary battery of metabolomics platforms was used, including untargeted metabolomics, Agilent dMRM Database and Method platform, and targeted metabolomics. Metabolite levels and RNA-metabolomics integrated modules were correlated with clinical parameters.

**Results:** Muscle samples from IBM patients had elevated TCA cycle intermediates with concomitant increase in anaplerotic amino acids, suggesting increased anaplerosis into the cycle. There was a decrease in upper glycolysis intermediates and an increase in most of the pentose phosphate pathway (PPP) metabolites. The PPP is the main source of NAPDH, a main antioxidant, and ribose-5-P a precursor of nucleic acids. There were marked sex-specific differences in the acylcarnitine profile, with a decrease in short-chain acylcarnitines only in males. Lastly, there was an increase in nucleic acid bases and a decrease in nucleotides. Several metabolites from various pathways had significant correlations with various clinical parameters, with the most pronounced sex-specific differences observed in correlations with acylcarnitines. RNA-metabolomics integration identified 4 modules, with the strongest correlation observed between one module and sex. The MMT score, an indicator of disease severity, showed a strong correlation with 3 modules. There were major sex specific differences with males having relatively similar correlation to the grouped (both sexes) analysis, while females had no significant correlation with any of the modules.

**Conclusion:** Taken together, our findings identified clinically significant alterations in central carbon metabolism in IBM, with major differences between males and females. Future studies are needed to determine the role of the detected metabolic alterations in IBM pathogenesis and track the changes longitudinally over the disease course.

## INTRODUCTION

Inclusion body myositis (IBM) is a disease of aging, characterized by progressive muscle loss, which eventually leads to loss of mobility.^1^ Skeletal muscles are a major consumer of energy and play a central role in metabolism.^2, 3^ Metabolomic studies on human muscle samples in myopathies are limited, partly due to the limited access to samples. IBM sits at the intersection of aging, chronic inflammation, and mitochondrial dysfunction, making it an ideal candidate for metabolomic studies.^4^ Despite being the most common muscle disease, only limited studies evaluated the metabolomic profile in IBM, and only one study performed on muscle tissue.^5^ In this study by Murakami et al, untargeted metabolomic analysis was performed on samples from 14 IBM patients, 10 males and 4 females, and 6 controls.^5^ The top 20 metabolites were associated with histamine biosynthesis, glycosaminoglycan metabolism, carnitine metabolism and creatine metabolism, and transcriptomics analysis on 5 samples was focused on the genes related to the identified metabolic pathways.^5^ In a study by Buzkova et al, 94 metabolites were evaluated in the sera of 6 IBM patients and patients with mitochondrial diseases and other neuromuscular disorders.^6^ IBM shared 39% of altered metabolites with progressive external ophthalmoplegia, a mitochondrial disease, versus only 23% with other neuromuscular disorders.^6^ Lastly, Canto-Santos et al evaluated organic acid levels in fibroblasts and urine samples, and nucleotides in urine samples from IBM patients and controls.^7^ 5 out of 19 organic acids alterations in fibroblasts reached statistical significance, with 2-hydroxyglutaric acid having the highest fold change. Only one organic acid (orotic acid) reached statistical significance in urine. Among nucleotides, orotidine and pseudo-uridine were significantly elevated. Furthermore, they evaluated metabolism-related genes in their RNA data and assessed their correlation with the observed changes in organic acids. Sex-specific differences in IBM metabolism have not been studied.

Our recently published study on a cohort of 38 IBM patients and 22 controls reported NLRP3 inflammasome activation and altered mitophagy in IBM.^8^ There was a significant elevation of p- S65-Ubiquitin, a mitophagy marker, in muscle lysates from male patients, whereas females had only mildly elevated levels that did not reach statistical significance.^8^ On histopathology, we demonstrated the accumulation of p-S65-Ubiquitin aggregates in cytochrome c oxidase (CCO) negative fibers that were strictly type II fibers. Indeed, CCO negative fibers in IBM muscles, with or without aggregates, were mainly type II.^8^ Type II fibers are more prone for mitochondrial dysfunction and have different metabolic signature than type I.^9^ Furthermore, the mitochondria play a central and multifaceted role in cellular metabolism, expanding beyond ATP synthesis to include the production and breakdown of metabolites, as well as the regulation of metabolic pathways through signaling.^10^ Furthermore, several metabolic reactions, including the tricarboxylic acid (TCA) cycle, occur inside the mitochondria. Hence, we aimed to investigate the metabolomic map of IBM, centered on the mitochondria, underscoring sex-specific changes.

## METHODS

### Ethical statement

The study was approved by the Mayo Clinic Institutional Review Board. The study was considered as a minimal risk; therefore, the requirement for informed consent was waived. However, the medical records of any patient who had not provided authorization for their use in research, in accordance with Minnesota statute 144.335, were not reviewed. The use of clinical residual muscle biopsy tissue was also approved by the Mayo Clinic Biospecimens Subcommittee.

### Study population

The study population consisted of 37 patients with IBM patients and 22 age and sex-matched controls without a myopathy. Patients with IBM fulfilled 2011 European Neuromuscular Centre (ENMC) diagnostic criteria for IBM.^11^ As patients had typical pattern of weakness and pathologically-confirmed diagnosis, they would also fulfill the recently published revision of those criteria.^12^ Controls consisted of patients without a myopathy, who underwent a muscle biopsy to rule out a muscle disorder and had an unremarkable work up. This is the same population that was included in our recently published study that included 38 patients with IBM and 22 controls.^8^ However, one IBM male patient had no sufficient leftover frozen muscle tissue to preform metabolomic analysis. All muscle biopsies were originally obtained for clinical purposes, and residuals were used for this project. Per our clinical practice standards, an open biopsy was performed in the operating room under conscious sedation after fasting overnight. The harvested muscle tissue was frozen in isopentane-cooled in liquid nitrogen and stored at - 80°C according to our Muscle laboratory protocols. About 30-50 mg of frozen tissue was used for the metabolomics experiments.

### Clinical parameters

Clinical parameters were obtained via chart review and included: age at biopsy, sex, disease duration (time from symptom onset till biopsy date), manual muscle testing (MMT) score and modified Rankin Scale (MRS). The MMT score is a summated motor exam of the bilateral shoulder abduction, elbow flexion, elbow extension, finger flexion, hip flexion, knee extension and ankle dorsiflexion. Each muscle is graded from 0 (normal) to 4 (complete paralysis), and the total score ranges from 0 (normal strength) to 56 (complete paralysis) as previously described.^13^ The MRS, used to assess patient’s level of disability, ranges from 0 (no symptoms) to 6 (dead) as follows: MRS 1- symptoms without disability; 2- slight disability with ability to perform baseline activities; 3- moderate disability requiring some help, but able to walk unassisted; 4- moderately severe disability with inability to walk unassisted (wheelchair bound); and 5- severe disability with need for constant nursing care and bedridden.^14^

### Metabolomics

Untargeted, Agilent dMRM Database and Method platform, and targeted metabolomics were performed. Detailed methods can be found in **Supplemental Methods**.

### Untargeted metabolomics

Untargeted metabolomics was performed as previously described.^15, 16^ Briefly, metabolites were extracted from frozen muscle samples and analyzed on a Quadrupole Time-of-Flight mass Spectrometer (MS) (Agilent Technologies 6550 Q-TOF) coupled with an Ultra High Pressure Liquid Chromatograph (Agilent Technologies 1290 Infinity UHPLC). Metabolites were profiled under both positive (p) and negative (n) electrospray ionization condition and separated using a hydrophilic interaction (hilic) and C18 columns (four modes: philic, nhilic, pC18 and nC18). Metabolites with log fold change > |0.58| (fold change >1.5) and adjusted *p-*value < 0.05 were considered significantly different between groups.

### Central Carbon metabolism

Agilent Metabolomics dMRM Database and Method platform focused on the central carbon metabolism was used (semi-targeted platform). Central carbon metabolites (219 compounds) were monitored and measured on muscle tissue homogenates using an Agilent 6460 triple quadrupole mass spectrometer couple with a 1290 Infinity II quaternary pump.^17^ Analytes were searched and confirmed against a curated dynamic multiple reaction monitoring (dMRM) database with retention time. Relative abundances between samples set are derived via multivariate analysis on Agilent Mass Profiler Professional software (MPP).

### Targeted metabolomics

The targeted metabolite panels performed on muscle tissue homogenates consisted of 11 TCA analytes measured by gas chromatography–MS (GC-MS), 10 glycolysis and pentose phosphate pathway (PPP) metabolites measured by GC-MS, 42 amino metabolites measured by liquid chromatography-mass spectrometry (LC-MS), carnitine and 13 acyl carnitines (specifically C0–C18:1) measured by LC-MS, and acetyl CoA measured by LC-MS. Metabolite measurements were normalized to the sample protein measurement by bicinchoninic acid-assay (BCA). Two group comparisons were performed via a 2-sample t test or a Wilcoxon rank sum test, as appropriate, using BlueSky statistics v.7.10 software and GraphPad Prism v.9.3.1.

### Transcriptomics

The raw gene-level count data from our recently published transcriptomics data from the same patient population were utilized for metabolomics-transcriptomics data integration.^8^

### Association between metabolite concentrations and clinical parameters

Central carbon metabolite data was normalized using trimmed means of M-values method and log2 transformed. Metabolites from targeted metabolomics and central carbon metabolism were pooled together. Correlation analysis with clinical parameters was performed using Person correlation.

### Integration of RNA-metabolomics data and correlation with clinical parameters

The gene expression count data was normalized using trimmed mean of M-values method and log2 transformed. Processed data was filtered to retain genes with >20% coefficient of variation (CV) of expression across the entire cohort. Filtered genes were z-score transformed.

Independently, the pooled metabolite data, described above) was filtered, like gene expression data using a CV filter of >20%, and z-score transformed. Scaled gene expression and metabolite data were combined for weighted co-expression network analysis.^18^ This process identifies modules, of genes and/or metabolites, that have strong association (by expression) with each other. Relationship between each module (via Eigen values) versus various continuous outcomes were measured using spearman correlation. Module-outcome correlation p-values were adjusted using Benjamini-Hochberg method. MetaboAnalyst 6.0 was used for joint pathway analysis in each module.^19^ Enrichment analysis was performed via a hypergeometric test, with degree centrality as topology measure and combine queries as integration methods. False discovery rate (FDR) q value of <0.05 was considered as statistically significant.

## RESULTS

### Study population

Patients’ demographics and disease characteristics are provided in **Table 1**. IBM group and controls had relatively similar age at biopsy. There were no significant differences in baseline characteristics between the two groups.

**TABLE 1:** Baseline characteristics and patient demographics.

| Variable | Inclusion body myositis<br>(n=37) | Controls (n=22) |
| --- | --- | --- |
| **Age at biopsy** | 62.7 (9.2) | 59.7 (10) |
| **Sex:** |  |  |
| Females | 19 (51) | 11 (50) |
| Males | 18 (49) | 11 (50) |
| **Race:** |  |  |
| American Indian/Alaskan Native | 1 (2.7) | 1 (4.5) |
| Black | 1 (2.7) | 0 |
| White | 31 (83.8) | 19 (86.4) |
| Unknown/not disclosed | 4 (10.8) | 2 (9.1) |
| **Biopsy target** |  |  |
| Quadriceps | 14 (37.9) | 8 (36.4) |
| Biceps brachii | 9 (24.3) | 5 (22.7) |
| Triceps brachii | 9 (24.3) | 3 (13.6) |
| Others | 5 (13.5) | 6 (27.3) |
| **Disease characteristics** |  |  |
| Disease duration at time of the biopsy (years) | 6.3 (3.9) | NA |
| Manual Motor Test score | 17 (9.3) | 0 |

Continuous variables are displayed as mean (SD) and categorical variables are displayed as the number of patients (percentage). NA: not applicable.

### Untargeted metabolomics

Results of untargeted metabolomics are shown in **(Figure 1)**. By nC18 mode, there were 36 metabolites that were significantly different between IBM samples and controls. Citric acid (TCA cycle) had the top variable importance in projection (VIP) score followed by 3-deoxyguanosine. D-Glycerate 2-phosphate (glycolysis) was among the top 15 metabolites. By pC18 mode, there were 63 metabolites that were significantly different between the two groups. Most of the metabolites with the top VIP scores consisted of nonendogenous or unidentified compounds.

**Figure 1:**
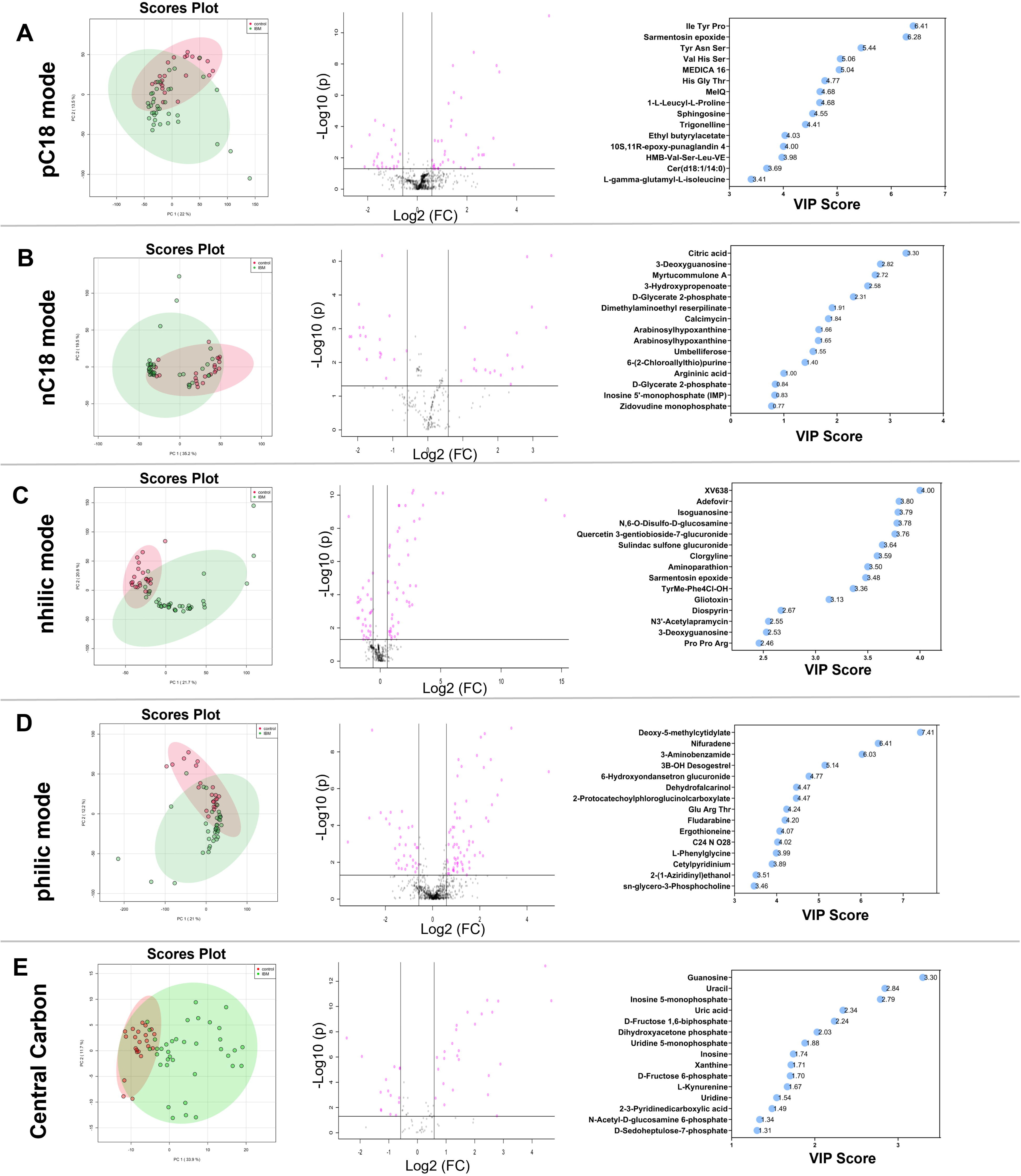
Results of untargeted metabolomics and central carbon metabolism. A) positive C18 (pC18) mode; B) negative C18 (nC18) mode; C) negative hydrophilic interaction (nhilic) mode, D) positive hydrophilic interaction (philic mode); E) central carbon metabolism. Each row displays primary component analysis scatter plot, volcano plot for changes in metabolites versus p values, and VIP scores.

Otherwise, list included many lipids and D-fructose which was elevated in IBM samples. By nhilic mode, there were several phospholipids and metabolites involved in eprenoid backbone synthesis which are essential to various cellular processes including membrane structure, electron transport, hormone synthesis and defense mechanisms. 3-Deoxyguanosine was again among metabolites with top VIP scores in addition to isoguanosine. By philic mode, metabolites with top VIP scores included several lipids. Analysis by sex showed relatively similar results.

### Central Carbon metabolism

With the fragmented nature of the results from the untargeted metabolomics, which is expected given the exploratory nature of the platform, we followed with the semi-targeted focused on the central carbon metabolism. It showed 41 metabolites that were significantly different between IBM samples and controls **(Figure 1)**. Metabolites with top VIP scores were mostly related to nucleic acids and glucose metabolism. IBM samples had elevated nucleic acid bases levels, but decreased monophosphate and diphosphate nucleotide levels compared to controls.

Furthermore, there was an increase in pentose phosphate pathway metabolites with a decrease in glycolysis pathway metabolites in IBM. Analysis by sex showed relatively similar results.

### Targeted metabolomics

Lastly, we homed in on the mitochondria-related metabolites by targeted panels. First, several TCA cycle metabolites were increased in IBM samples with no difference in lactate level between IBM and controls **(Figure 2)**. The changes in some of the glycolysis metabolites suggest a decrease in upper glycolysis. Simultaneously, most of the PPP metabolites were increased in IBM indicating a shift to PPP **(Figure 3)**. PPP is one of the main pathways that generate NADPH, involved in suppressing oxidative stress, and ribose-5-phosphate, used for nucleotide synthesis.^20^ These results were relatively similar between males and females.

**Figure 2:**
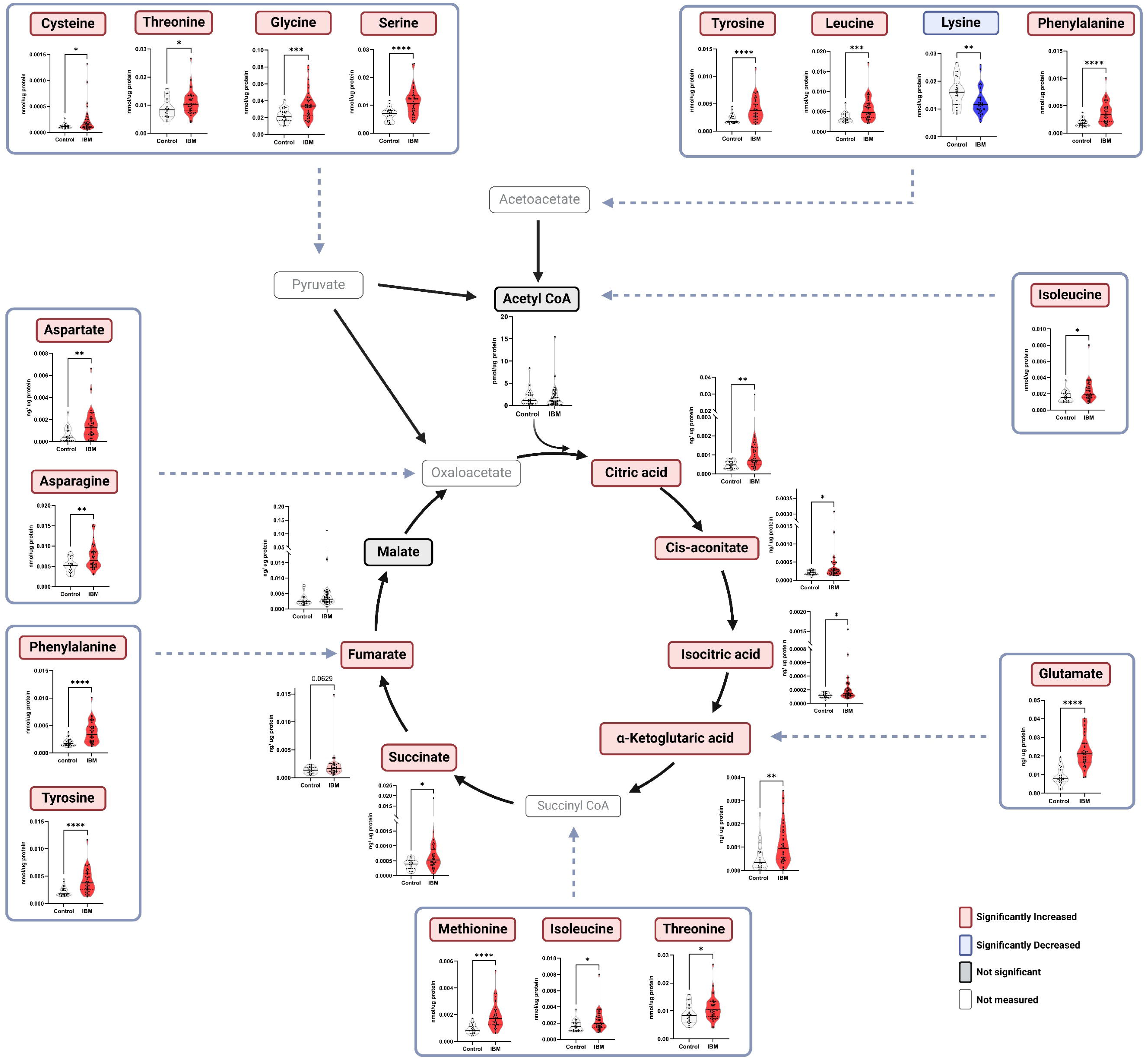
Targeted metabolomics results for tricarboxylic acid (TCA) cycle and anaplerotic amino acids levels. Individual data are presented as violin plots.

**Figure 3:**
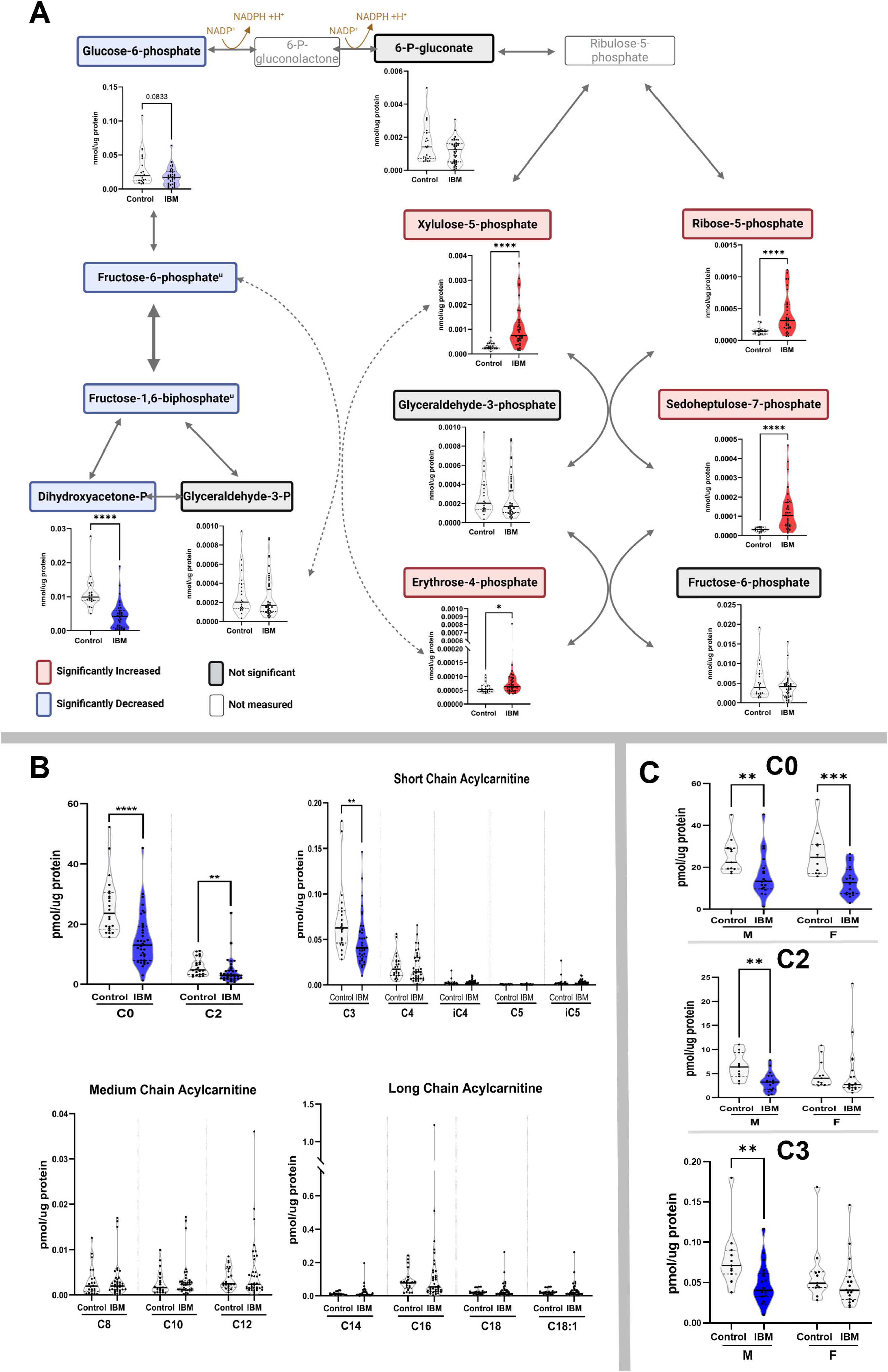
Upper glycolysis-pentose phosphate pathway and acylcarnitine profile. A) Upper glycolysis and pentose phosphate pathway metabolites. Individual data for targeted metabolomics are shown as violin plots; u: metabolites from central carbon metabolism platform. B&C) Acylcarnitine profile; results by sex shown in (C). Acylcarnitines are displayed according to the number of carbon atoms.

On the acylcarnitine profile, carnitine and short-chain chain acylcarnitine levels (C2, C3) were decreased in IBM **(Figure 3)**. The decrease in short chain acylcarnitine levels was only observed in males. There was no significant difference in acetyl-CoA levels between IBM and controls.

On the amino acid profile, 22 out of 42 amino acids were altered in IBM. Notably, there an anaplerotic increase in several amino acids serving the TCA cycle **(Figure 2)**.

### Association between metabolite concentrations and clinical parameters

Results from the correlation analysis between targeted metabolites and some of the central carbon metabolism platform metabolites are shown in **(Figure 4).** The MMT score had a significant positive correlation with 13 metabolites. The metabolite with the strongest positive correlation (higher levels associated with more severe weakness) was cysteine. Cysteine is the rate-limiting substrate of the synthesis of glutathione, an antioxidant that neutralizes reactive oxygen species.^21^ A PPP metabolite (sedoheptulose-7-P) and a glycolysis inhibitor (2-deoxy-D-glucose-6-P) positively correlated with MMT, whereas glycolysis metabolites (glucose-6-P, fructose-6-P and glycerate) inversely correlated with MMT. Carnosine and sarcosine were inversely correlated with MMT. Sarcosine, a derivative of creatine metabolism, is an autophagy activator modulated by aging and is essential for muscle health.^22, 23^ Carnosine, formed by β-alanine and L-histidine, is involved in various roles during exercise and increasing intramuscular carnosine is being considered a therapeutic target to enhance exercise performance.^24^ Decreased Taurine levels in blood have been associated with aging. ^25^ However, in IBM muscle samples, higher levels of Taurine were (positively) correlated with higher MMT. MMT was correlated with the levels of nucleic acid bases and inversely correlated with monophosphate and diphosphate nucleotides. Only 9 metabolites correlated with MRS. Most were nucleic acids, in addition to L-Cystathione and S-5-Adenosyl-L-homocyteine (related to antioxidation). Cyclic GMP (cGMP) inversely correlated with MRS. Decreased cGMP levels have been associated with aging and aging related disorders, and here, decreased cGMP levels were associated with higher disability (higher MRS).^26^ Only alpha-aminoadipic acid (positively) correlated with disease duration. Alpha-aminoadipic acid is considered a marker for glucose metabolism and diabetes and has a potential role in glutamatergic neurotransmission and oxidative stress.^27, 28^ Four metabolites inversely correlated with disease duration, including creatinine. Age at biopsy (positively) correlated with 3 TCA cycle metabolites and inversely with thiamine among others.

**Figure 4:**
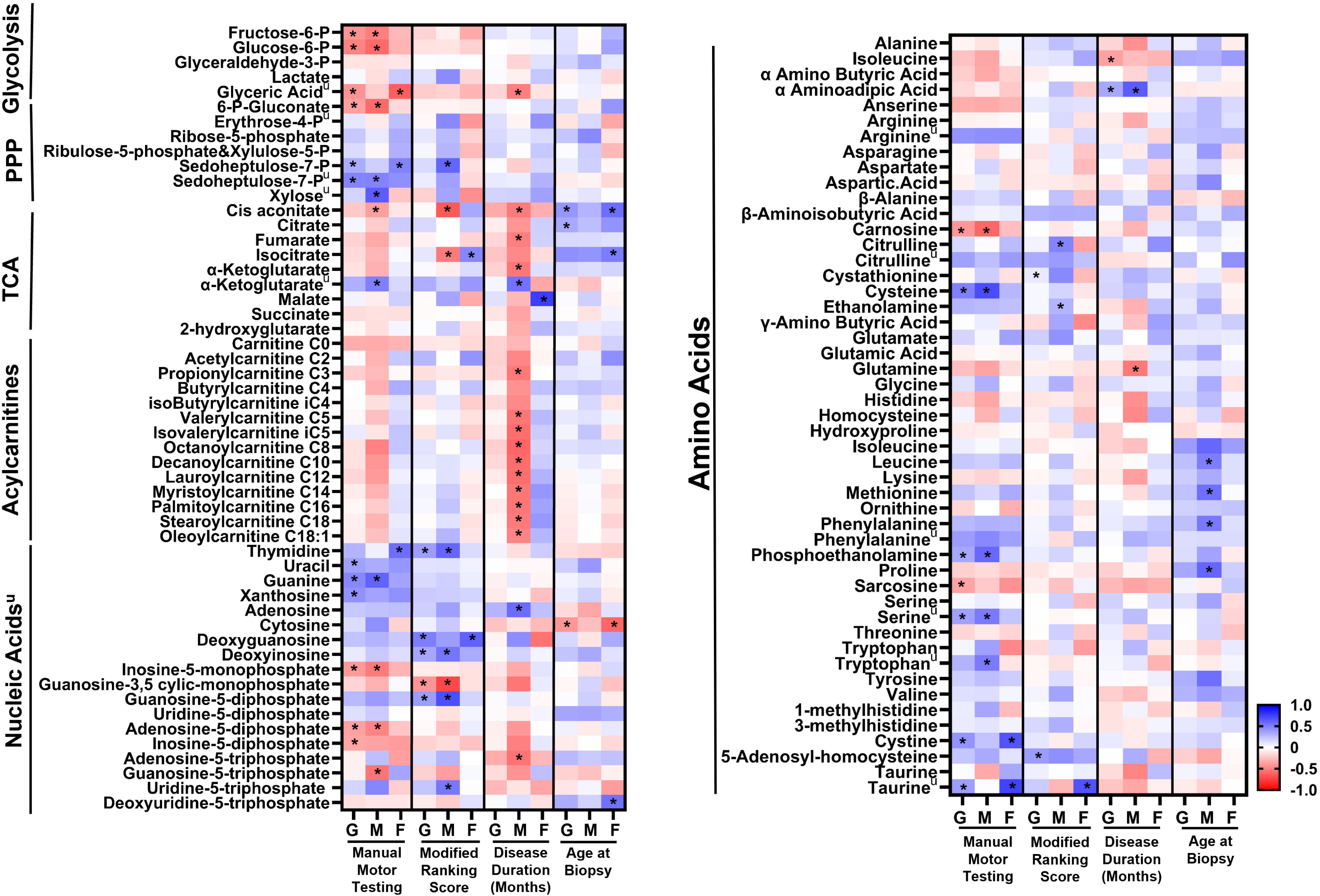
Correlation of metabolites with clinical parameters. Correlations that reach statistical significance are marked with a (*). u: metabolites from central carbon metabolism platform.

When analyzing by sex, there were sex specific differences. For MMT, males had shared metabolites with the grouped (both sexes) comparison related to PPP, glycolysis and nucleosides. Cysteine had the strongest correlation with MMT in males. In contrast, MMT in females had the strongest correlation with Taurine and there were less correlations that reached statistical significance in females. Males and females shared only one common metabolite that correlated with MMT: 3-(2-hydroxyethyl)indole that had opposite correlation: positive in males and negative in females. Its role in human biology is unclear. For MRS, similar pattern is observed with no common metabolites between males and females. Again, Taurine, in females, had the strongest correlation with MRS. The most striking sex specific differences were in the metabolomic correlates of disease duration. Similar to the group analysis, a small number of metabolites had significant correlation in females. However, 30 metabolites had significant correlations in males including 10 acylcarnitines and TCA cycle intermediates, all of which had inverse correlation (higher levels with shorter disease duration).

### RNA-Metabolomics integration and correlation with clinical parameters

Unsupervised integration of bulk RNA sequencing data with metabolites from the targeted analysis and central carbon metabolism platform yielded 9 modules **(Figure 5**). Correlation with clinical parameters showed sex had the strongest (inverse) correlation with the Pink module.

**Figure 5:**
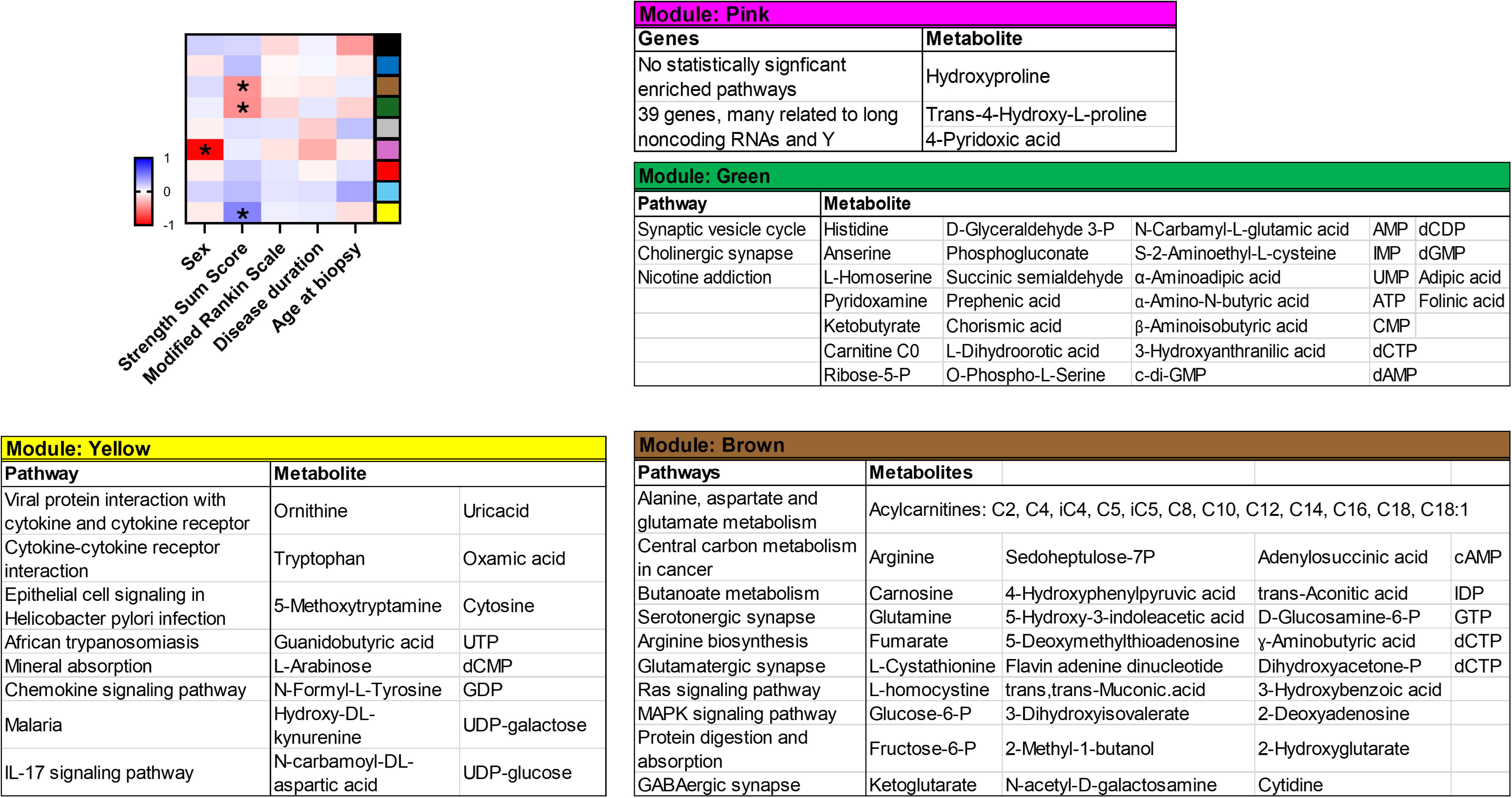
Correlation of clinical parameters with RNA-metabolites integrated modules. Correlations that reached statistical significance are marked with a (*) and corresponding modules displayed as tables.

This was followed by the MMT, reflecting disease severity, that correlated with the Yellow module and inversely correlated with the Brown and Green modules. Age at biopsy, disease duration and MRS had no significant correlation with any of the modules. The Pink Module contained 3 metabolites. Hydroxyproline is a critical component of collagen and essential for muscle elasticity and repair; trans-4-Hydroxy L-proline is the isomer found in collagen.^29^ Pyridoxic acid (vitamin B6) is essential for glycogenolysis and muscle contraction, and vitamin B6 deficiency has been associated with sarcopenia.^30^ Although no particular pathway was enriched in the 39 genes of the Pink Module, there were several long noncoding RNAs and Y chromosome genes. Among these genes, UTY is thought to contribute to sex-specific phenotypes and disease risks.^31^ In addition, there were several genes related to regulation of gene expression, protein homeostasis, vesicular trafficking and NADPH. The Yellow module included 15 metabolites and 351 genes **(Figure 5).** Metabolites included 3 related to tryptophan metabolism, ornithine and 5 nucleic acids among others. Tryptophan is a precursor of serotonin and vitamin B3 and is important for skeletal muscle energy production and metabolism. It is essential for the kynurenine pathway that is related to sarcopenia, aging and aging-related inflammation.^32^ Ornithine, a nitrogenous amino acid, is important for nitrogen metabolism and counteracting fatigue.^33^ Enriched pathways were related to cytokines/chemokines, IL-17 signaling and neuroactive-ligand-receptor interaction. Many of the listed pathways had overlapping genes related to cytokines and chemokine receptors: *CXCR1, CXCR2, CXCL1, CXCL8* (IL8)*, PF4* (a.k.a *CXCL4*)*, IL6, CCL21*. In addition to the listed pathways, we note 4 genes were related to the NLRP inflammasomes (*NLRP12, MEFV, CCL21 & CXCL8*) and *GDF15*. GDF15 is a myokine, considered a mitochondrial marker and a diagnostic biomarker for IBM and mitochondrial disorders.^34^ In the Green Module, there were 31 metabolites and 195 genes. Metabolites included: carnitine, beta-aminoisobutyric acid, alpha-aminoadipic acid, 3 related to PPP, 2 to vitamin B6 metabolism, 3 to histidine metabolism, and 9 nucleic acids among others. Beta-aminoisobutyric acid is produced by skeletal muscle during exercise and acts like a myokine that regulates metabolism, oxidative stress and inflammation.^35^ Similarly, pathways enriched in the Green module were mostly related to neurotransmission. Lastly, the Brown Module had 44 metabolites and 369 genes. Metabolites included several related to glycolysis, TCA, PPP, nucleic acids, acylcarnitines in addition to carnosine, L-cystathionine and others. Enriched pathways were related to central carbon metabolism, MAPK signaling, RAS signaling, neurotransmission and others. MAPK and RAS signaling pathways are regulatory pathways linked to cell growth, survival and response to oxidative stress, invoked in neurodegenerative diseases and being considered as therapeutic targets in these disorders.^36, 37^

Next, we correlated the clinical parameters with the modules from RNA-metabolomics integration for each sex separately. In females, none of the modules had a significant correlated with any of the clinical parameters. In males, 3 modules correlated with MMT with many shared metabolites and pathways with the grouped analysis. The Black Module shared similar enriched pathways with the Yellow Module, and the Red Module with the Green Module.

## DISCUSSION

In this study, we provide a mitochondria-centered comprehensive metabolomic map of the skeletal muscle of IBM via various platforms, underscoring common and sex-specific metabolic changes. We also correlate the findings with clinical parameters to determine their clinical significance. Lastly, we perform RNA-metabolomic integration and identify modules of genes and metabolites that are associated with various clinical parameters.

Majority of TCA cycle intermediates were elevated in skeletal muscles of IBM patients, without any changes in lactate or acetyl-CoA levels. There are limited studies evaluating TCA metabolites in human muscle tissue in general and in muscle disease in particular, as most studies were pursued in cellular or animal models.^38, 39^ In a study evaluating sarcopenia in patients with hip fracture, serum form those patients had elevated TCA metabolites.^40^ The pattern was different compared to the one seen in our samples (decreased α-increased malate, fumarate and succinate) and the authors postulated it was related to complex I deficiency and subsequent buildup of NADH. An increase in TCA metabolites has also been reported in the postmortem brains of patients with Alzheimer’s disease.^41^ In our study, many of TCA cycle-related amino acids (**Figure 2**) were increased, suggesting increased anaplerosis in IBM that should be further explored in future studies. In a study by Motori et al., anaplerosis was reported to play a role in offsetting oxidative phosphorylation defects in neurodegenerative diseases.^42^ Increased anaplerosis has also been observed in other disorders such as cancer.^43^

Lastly, TCA metabolites had significant correlations with clinical parameters, with remarkable sex-specific changes, indicating the findings are of clinical significance. The most consistent pattern was seen with disease duration correlation, only in males, where higher levels of all TCA metabolites were associated with shorter disease duration. Correlation with age at biopsy was significant in females where younger patients had higher TCA levels. From the integrated RNA- metabolites modules, the Brown module, which correlated with MMT score, contained several TCA cycle intermediates. With the increase in several central carbon metabolites, it remains unclear whether a hypermetabolic state exists in IBM. The “energy constraint model” is a recently proposed innovative concept, where underlying defects in mitochondrial oxidative phosphorylation drive an energetically costly hypermetabolic state that is detrimental to the cell and could cause accelerated aging.^44^

The other main finding in our study was the decrease in upper glycolysis metabolites, including glucose-6-P and Fructose-6-P, and an increase in most PPP metabolites, suggestive of a shift from glycolysis to PPP. The shift to PPP has two potential reasons: generation of NADPH to defeat oxidative stress, and synthesis of nucleotides through ribose-5-P.^45^ Cysteine, the rate limiting substrate for glutathione synthesis had the strongest correlation with MMT and nucleic acid metabolism was markedly perturbed. Activation of PPP is thought to play a protective role in neurodegenerative diseases.^46^ Interestingly, abnormal glucose metabolism has also been linked to other diseases of aging, including Alzheimer’s disease, where decreased glucose metabolism on 18-F fluorodeoxyglucose positron emission tomography is one of the first manifestations of this disorder.^47^ Studies on PPP in human skeletal muscle are rare and to the best of our knowledge, a shift to PPP has not been addressed in a musculoskeletal disease.

The decrease in glucose-6-P and fructose-6-P was associated with higher MMT scores indicating more severe weakness. Although it did not reach statistical significance, the pattern was consistent for the majority of the metabolites with decreased levels of glycolysis metabolites and increased levels of PPP metabolites associated with more severe MMT score. Likewise, the RNA-metabolites modules that correlated with MMT contained several PPP and glycolysis metabolites.

The main sex-specific difference in the levels of metabolites between males and females was observed in the acylcarnitine profile where the grouped and the male only analyses showed a decrease in short chain acylcarnitines (C0, C2 and C3), whereas females had only decreased carnitine levels. The sex differences in acylcarnitine metabolism were striking in the correlation analysis as there was a strong inverse correlation, only in males, between disease duration and most acylcarnitine levels. Although it didn’t reach statistical significance, almost all acylcarnitines had inverse correlation with MMT in males and positive (but weak) in females. Likewise, the Brown module included several acylcarnitines. These sex-specific alterations in lipid metabolism are intriguing and require further investigation.

Several amino acids were altered in IBM samples, many of which related to skeletal muscle health and aging, such as sarcosine and carnosine, as discussed in the results section. Additionally, branched chain amino acids (BCAA) leucine and isoleucine were elevated in IBM. Increased BCAAs are thought to promote oxidative stress and inflammation in human mononuclear cells, by mTORC1 activation.^48^ Decreased taurine levels have been associated with aging, whereas elevated levels, as in our samples, were suggested to help alleviating the oxidative stress and sarcopenia.^25, 49, 50^ Serine, methionine and sarcosine are connected through the one-carbon cycle, disturbances of which have been closely linked to mitochondrial dysfunction.^51^

As captured by central carbon metabolism platform, there were remarkable nucleic acid alterations with an increase in nucleic acid bases and a decrease in mono- and diphosphate nucleotides. The reason for these changes is unclear and they may indicate increased purine and pyrimidine breakdown or decreased synthesis. It could also indicate disrupted cellular energy metabolism, where the high demand for ATP is not being met. Decreased electron transport chain activity has been linked to activation of PPP, decreased de novo purine synthesis, and increased uptake of extracellular bases.^52^ The central carbon metabolism platforms also revealed several altered lipids that require further investigation via a lipidomic platform.

There was a consistent pattern when evaluating for sex-specific changes: in general, males consistently had several correlations with clinical parameters whether with metabolites only or RNA-metabolites integrated modules. In contrast, females had fewer metabolites and none of the RNA-metabolites integrated modules correlating with clinical parameters. The reason for that remains elusive. It could be that females are more resilient to metabolic changes caused by the disease. It could also be that the interaction between metabolic disturbances and clinical parameters is more variable in females so that none is significant when evaluated on a group level. Of note, the Pink Module which had the strongest correlation with a clinical parameter: biologic sex, consisted of several long noncoding RNAs and Y chromosome-related genes. Understanding the regulator role of noncoding RNAs is a growing field.^53^ The role of gonosomal genes in metabolic regulation in musculoskeletal and neurodegenerative diseases is not well explored.

Our cross-sectional study provides a comprehensive evaluation of central carbon metabolism in IBM performed on frozen samples obtained from IBM patients. Inherent to this design and the nature of the specimens, we cannot determine which alterations are detrimental versus adaptive and we cannot perform interventions of the specimens to further explore pathomechanisms or determine fluxes. IBM does not have a valid disease model, as the commonly used mouse models in the literature are deemed not representative of the disease.^12, 54^ On the other hand, especially in metabolomics, performing the experiments on samples derived from patients offers a direct window into disease pathogenesis which would facilitate bench to bedside translation.^54^

Furthermore, if diagnostic or prognostic biomarkers are to arise from these studies, this could be directly applied to samples obtained in clinical practice. Of note, correlation analysis does not imply a causative relationship between the identified metabolomic abnormalities and the clinical parameters. Additional limitations include the retrospective nature of the chart review limiting the clinical parameters that can be abstracted. Furthermore, the variability in the muscles that were biopsied, patients’ medications at time of the biopsy, and the quality of the tissue could have contributed. However, IBM does not have any known effective treatment, the variability in muscle biopsies was in both groups and the samples were obtained and handled in the same method in both groups, which would help offsetting these limitations to some extent. Unlike *in-vivo* and *in-vitro* studies, there will always be variability in the conditions of the samples obtained from patients.

Future studies are needed to further evaluate the role of the detected metabolic alterations in IBM pathogenesis and to further explore the various metabolites identified on the untargeted metabolomics platforms. Furthermore, assessing the longitudinal changes in metabolites over the disease course would be of interest.

## ACKNOWLEDGEMENTS

This research was supported by grants from the National Institute of Arthritis, Musculoskeletal and Skins Diseases (NIAMS, K08-AR78254), American Neuromuscular Foundation (ANF #2, career development grant), and Mayo Clinic Center for Clinical and Translational Sciences (small project grant and Team Science pilot award, UL1TR002377 from the National Center for Advancing Translational Sciences) to E.N. The study contents are solely the responsibility of the authors and do not necessarily represent the official view of the NIH and other funding organizations. The funders had no role in the study design, data collection and analysis, the decision to publish, or the preparation of the manuscript.

## REFERENCES

1. Naddaf E, Shelly S, Mandrekar J, et al. Survival and associated comorbidities in inclusion body myositis. Rheumatology (Oxford, England) 2022;61:2016–2024.

2. Wolfe RR. The underappreciated role of muscle in health and disease. Am J Clin Nutr 2006;84:475–482.

3. Zurlo F, Larson K, Bogardus C, Ravussin E. Skeletal muscle metabolism is a major determinant of resting energy expenditure. J Clin Invest 1990;86:1423–1427.

4. Guglielmi V, Cheli M, Tonin P, Vattemi G. Sporadic Inclusion Body Myositis at the Crossroads between Muscle Degeneration, Inflammation, and Aging. Int J Mol Sci 2024;25.

5. Murakami A, Noda S, Kazuta T, et al. Metabolome and transcriptome analysis on muscle of sporadic inclusion body myositis. Ann Clin Transl Neurol 2022;9:1602–1615.

6. Buzkova J, Nikkanen J, Ahola S, et al. Metabolomes of mitochondrial diseases and inclusion body myositis patients: treatment targets and biomarkers. EMBO Mol Med 2018;10.

7. Cantó-Santos J, Valls-Roca L, Tobías E, et al. Unravelling inclusion body myositis using a patient-derived fibroblast model. J Cachexia Sarcopenia Muscle 2023;14:964–977.

8. Naddaf E, Nguyen TKO, Watzlawik JO, et al. NLRP3 inflammasome activation and altered mitophagy are key pathways in inclusion body myositis. medRxiv 2024:2024.2006.2015.24308845.

9. Wischnewski S, Thäwel T, Ikenaga C, et al. Cell type mapping of inflammatory muscle diseases highlights selective myofiber vulnerability in inclusion body myositis. Nat Aging 2024;4:969–983.

10. Spinelli JB, Haigis MC. The multifaceted contributions of mitochondria to cellular metabolism. Nat Cell Biol 2018;20:745–754.

11. Rose MR. 188th ENMC International Workshop: Inclusion Body Myositis, 2-4 December 2011, Naarden, The Netherlands. Neuromuscul Disord 2013;23:1044–1055.

12. Lilleker JB, Naddaf E, Saris CGJ, Schmidt J, de Visser M, Weihl CC. 272nd ENMC international workshop: 10 Years of progress - revision of the ENMC 2013 diagnostic criteria for inclusion body myositis and clinical trial readiness. 16-18 June 2023, Hoofddorp, The Netherlands. Neuromuscul Disord 2024;37:36–51.

13. Pinto MV, Laughlin RS, Klein CJ, Mandrekar J, Naddaf E. Inclusion body myositis: correlation of clinical outcomes with histopathology, electromyography and laboratory findings. Rheumatology (Oxford, England) 2022;61:2504–2511.

14. van Swieten JC, Koudstaal PJ, Visser MC, Schouten HJ, van Gijn J. Interobserver agreement for the assessment of handicap in stroke patients. Stroke 1988;19:604–607.

15. Trushina E, Dutta T, Persson XM, Mielke MM, Petersen RC. Identification of altered metabolic pathways in plasma and CSF in mild cognitive impairment and Alzheimer’s disease using metabolomics. PloS one 2013;8:e63644.

16. Dutta T, Chai HS, Ward LE, et al. Concordance of changes in metabolic pathways based on plasma metabolomics and skeletal muscle transcriptomics in type 1 diabetes. Diabetes 2012;61:1004–1016.

17. Lee H-J, Kremer DM, Sajjakulnukit P, Zhang L, Lyssiotis CA. A large-scale analysis of targeted metabolomics data from heterogeneous biological samples provides insights into metabolite dynamics. Metabolomics : Official journal of the Metabolomic Society 2019;15:103.

18. Langfelder P, Horvath S. WGCNA: an R package for weighted correlation network analysis. BMC bioinformatics 2008;9:559.

19. Pang Z, Lu Y, Zhou G, et al. MetaboAnalyst 6.0: towards a unified platform for metabolomics data processing, analysis and interpretation. Nucleic acids research 2024;52:W398–w406.

20. TeSlaa T, Ralser M, Fan J, Rabinowitz JD. The pentose phosphate pathway in health and disease. Nat Metab 2023;5:1275–1289.

21. Papet I, Rémond D, Dardevet D, et al. Chapter 21 - Sulfur Amino Acids and Skeletal Muscle. In: Walrand S, ed. Nutrition and Skeletal Muscle: Academic Press, 2019: 335–363.

22. Walters RO, Arias E, Diaz A, et al. Sarcosine Is Uniquely Modulated by Aging and Dietary Restriction in Rodents and Humans. Cell Reports 2018;25:663–676.e666.

23. Gu X, Wang W, Yang Y, et al. The Effect of Metabolites on Mitochondrial Functions in the Pathogenesis of Skeletal Muscle Aging. Clin Interv Aging 2022;17:1275–1295.

24. Perim P, Marticorena FM, Ribeiro F, et al. Can the Skeletal Muscle Carnosine Response to Beta-Alanine Supplementation Be Optimized? Frontiers in Nutrition 2019;6.

25. Singh P, Gollapalli K, Mangiola S, et al. Taurine deficiency as a driver of aging. Science (New York, NY) 2023;380:eabn9257.

26. Kelly MP. Cyclic nucleotide signaling changes associated with normal aging and age-related diseases of the brain. Cellular signalling 2018;42:281–291.

27. Wang TJ, Ngo D, Psychogios N, et al. 2-Aminoadipic acid is a biomarker for diabetes risk. J Clin Invest 2013;123:4309–4317.

28. da Silva JC, Amaral AU, Cecatto C, et al. α-Aminoadipic Acid Cause Disturbance of Glutamatergic Neurotransmission and Induction of Oxidative Stress In Vitro in Brain of Adolescent Rats. Neurotox Res 2017;32:276–290.

29. Xu S, Gu M, Wu K, Li G. Unraveling the Role of Hydroxyproline in Maintaining the Thermal Stability of the Collagen Triple Helix Structure Using Simulation. J Phys Chem B 2019;123:7754–7763.

30. Kato N, Kimoto A, Zhang P, et al. Relationship of Low Vitamin B6 Status with Sarcopenia, Frailty, and Mortality: A Narrative Review. Nutrients 2024;16:177.

31. Rock KD, Folts LM, Zierden HC, et al. Developmental transcriptomic patterns can be altered by transgenic overexpression of Uty. Scientific Reports 2023;13:21082.

32. Ballesteros J, Rivas D, Duque G. The Role of the Kynurenine Pathway in the Pathophysiology of Frailty, Sarcopenia, and Osteoporosis. Nutrients 2023;15.

33. Sahu B, Pani S, Swalsingh G, et al. Long-term physical inactivity induces significant changes in biochemical pathways related to metabolism of proteins and glycerophospholipids in mice. Mol Omics 2024;20:64–77.

34. De Paepe B, Verhamme F, De Bleecker JL. The myokine GDF-15 is a potential biomarker for myositis and associates with the protein aggregates of sporadic inclusion body myositis. Cytokine 2020;127:154966.

35. Yi X, Yang Y, Li T, et al. Signaling metabolite β-aminoisobutyric acid as a metabolic regulator, biomarker, and potential exercise pill. Frontiers in endocrinology 2023;14:1192458.

36. Ahmed T, Zulfiqar A, Arguelles S, et al. Map kinase signaling as therapeutic target for neurodegeneration. Pharmacological Research 2020;160:105090.

37. Gravandi MM, Abdian S, Tahvilian M, et al. Therapeutic targeting of Ras/Raf/MAPK pathway by natural products: A systematic and mechanistic approach for neurodegeneration. Phytomedicine 2023;115:154821.

38. Li H, Uittenbogaard M, Navarro R, et al. Integrated proteomic and metabolomic analyses of the mitochondrial neurodegenerative disease MELAS. Mol Omics 2022;18:196–205.

39. Garvey SM, Dugle JE, Kennedy AD, et al. Metabolomic profiling reveals severe skeletal muscle group-specific perturbations of metabolism in aged FBN rats. Biogerontology 2014;15:217–232.

40. Hintze S, Baber L, Hofmeister F, et al. Exploration of mitochondrial defects in sarcopenic hip fracture patients. Heliyon 2022;8:e11143.

41. Kurano M, Saito Y, Yatomi Y. Comprehensive Analysis of Metabolites in Postmortem Brains of Patients with Alzheimer’s Disease. Journal of Alzheimer’s disease : JAD 2024;97:1139–1159.

42. Motori E, Atanassov I, Kochan SMV, et al. Neuronal metabolic rewiring promotes resilience to neurodegeneration caused by mitochondrial dysfunction. Sci Adv 2020;6:eaba8271.

43. Gonsalves WI, Ramakrishnan V, Hitosugi T, et al. Glutamine-derived 2-hydroxyglutarate is associated with disease progression in plasma cell malignancies. JCI Insight 2018;3.

44. Sercel AJ, Sturm G, Gallagher D, et al. Hypermetabolism and energetic constraints in mitochondrial disorders. Nat Metab 2024;6:192–195.

45. Stincone A, Prigione A, Cramer T, et al. The return of metabolism: biochemistry and physiology of the pentose phosphate pathway. Biol Rev Camb Philos Soc 2015;90:927–963.

46. Tang BL. Neuroprotection by glucose-6-phosphate dehydrogenase and the pentose phosphate pathway. J Cell Biochem 2019;120:14285–14295.

47. Chang R, Trushina E, Zhu K, et al. Predictive metabolic networks reveal sex- and APOE genotype-specific metabolic signatures and drivers for precision medicine in Alzheimer’s disease. Alzheimers Dement 2023;19:518–531.

48. Zhenyukh O, Civantos E, Ruiz-Ortega M, et al. High concentration of branched-chain amino acids promotes oxidative stress, inflammation and migration of human peripheral blood mononuclear cells via mTORC1 activation. Free Radic Biol Med 2017;104:165–177.

49. Lambert IH, Kristensen DM, Holm JB, Mortensen OH. Physiological role of taurine--from organism to organelle. Acta Physiol (Oxf) 2015;213:191–212.

50. Scicchitano BM, Sica G. The Beneficial Effects of Taurine to Counteract Sarcopenia. Curr Protein Pept Sci 2018;19:673–680.

51. Bao XR, Ong SE, Goldberger O, et al. Mitochondrial dysfunction remodels one-carbon metabolism in human cells. Elife 2016;5.

52. Wu Z, Bezwada D, Cai F, et al. Electron transport chain inhibition increases cellular dependence on purine transport and salvage. Cell metabolism 2024;36:1504–1520.e1509.

53. Statello L, Guo C-J, Chen L-L, Huarte M. Gene regulation by long non-coding RNAs and its biological functions. Nature Reviews Molecular Cell Biology 2021;22:96–118.

54. Skolka MP, Naddaf E. Exploring challenges in the management and treatment of inclusion body myositis. Current opinion in rheumatology 2023;35:404–413.

55. Dutta T, Kudva YC, Persson XM, et al. Impact of Long-Term Poor and Good Glycemic Control on Metabolomics Alterations in Type 1 Diabetic People. The Journal of clinical endocrinology and metabolism 2016;101:1023–1033.

56. Wilkins J, Sakrikar D, Petterson XM, Lanza IR, Trushina E. A comprehensive protocol for multiplatform metabolomics analysis in patient-derived skin fibroblasts. Metabolomics : Official journal of the Metabolomic Society 2019;15:83.

57. Cipollina C, ten Pierick A, Canelas AB, et al. A comprehensive method for the quantification of the non-oxidative pentose phosphate pathway intermediates in Saccharomyces cerevisiae by GC-IDMS. J Chromatogr B Analyt Technol Biomed Life Sci 2009;877:3231–3236.

58. Lanza IR, Zhang S, Ward LE, Karakelides H, Raftery D, Nair KS. Quantitative Metabolomics by 1H-NMR and LC-MS/MS Confirms Altered Metabolic Pathways in Diabetes. PloS one 2010;5:e10538.

59. Gonsalves WI, Broniowska K, Jessen E, et al. Metabolomic and Lipidomic Profiling of Bone Marrow Plasma Differentiates Patients with Monoclonal Gammopathy of Undetermined Significance from Multiple Myeloma. Scientific Reports 2020;10:10250.

