## Supplemental Methods for "Mitochondria-centered metabolomic map of inclusion body myositis: sex-specific alterations in central carbon metabolism"

**Untargeted metabolomics:**
Pulverized tissue was homogenized via sonication after adding 1 x PBS (10mL per mg tissue) prior to prepping. Tissue homogenates were deproteinized with six times volume of cold acetonitrile:methanol (1:1 ratio) after the addition of ^13^C_6_-phenylalanine (5 µl at 100ng/µl) as internal standard, kept on ice with intermittent vortexing then centrifuged at 18000xg for 30 minutes at 4C. The supernatants were divided into 4 aliquots and dried down using a stream of nitrogen gas for analysis on a Quadruple Time-of-Flight Mass Spectrometer (Agilent Technologies 6550 Q-TOF) coupled with an Ultra High Pressure Liquid Chromatograph (Agilent Technologies 1290 Infinity UHPLC). Profiling data was acquired under both positive and negative electrospray ionization conditions over a mass range m/z of 100 - 1200 at a resolution of 10,000 (separate runs) in scan mode. Metabolite separation was achieved using two columns of different polarity, a hydrophilic interaction column (Waters HILIC, ethylene-bridged hybrid 2.1 x 150 mm, 1.7 mm) and a reversed-phase C18 column (Waters high-strength silica 2.1 x 150 mm, 1.8 mm) with gradient described previously.^15, 16, 55^ Run time was 18 min for HILIC and 27 min for C18 column using a flow rate of 400µl/min. A total of four runs per sample were performed to give maximum coverage of metabolites. Samples were injected in duplicate or triplicate, wherever necessary, and a pooled quality control (QC) sample, made up of all the samples from each study was injected several times during a run. A separate plasma quality control (QC) sample was analyzed with pooled QC to account for analytical and instrumental variability. Dried samples were stored at -20^o^C until analysis. Samples were reconstituted in running buffer and analyzed within 48 hours of reconstitution. Auto-MS/MS data was also acquired with pooled QC sample to aid in unknown compound identification using fragmentation pattern.

Data analysis: Data alignment, filtering, univariate, multivariate statistical and differential analysis was performed using Mass Profiler Professional (Agilent Inc, USA). Metabolites detected in at least ≥80% of one of two groups were selected for differential expression analyses. Metabolite peak intensities and differential regulation of metabolites between groups were determined as described previously. ^15, 16, 55^ Each sample was normalized to the internal standard and log 2 transformed. Unpaired t-test with multiple testing correction p<0.05 was used to find the differentially expressed metabolites between two groups. Default settings were used with the exception of signal-to-noise ratio threshold (3), mass limit (0.0025 units), and time limit (9 s). Putative identification of each metabolite was done based on accurate mass (m/z) against METLIN database using a detection window of ≤5 ppm. The putatively identified metabolites were annotated as Chemical Abstracts Service (CAS), Kyoto Encyclopedia of Genes and Genomes (KEGG), Human Metabolome Project (HMP) database, and LIPID MAPS identifiers (LM_ID).

**Central carbon metabolism**

Muscle tissue homogenates were deproteinized by adding 500ul chilled methanol and acetonitrile solution to cell lysates, followed by lipids removal on Agilent Captiva ND-lipids plates. Cleaned samples were dried down and brought up in running buffer prior to LCMS analysis. Central carbon metabolites (219 compounds) were monitored and measured on an Agilent 6460 triple quadrupole mass spectrometer couple with a 1290 Infinity II quaternary pump. Acquisition was captured in negative electrospray ionization and dynamic multiple reaction monitoring (dMRM) post ion-pairing reverse phase chromatographic separation. Analytes were searched and confirmed against a curated dMRM database with retention time. Relative abundances between samples set are derived via multivariate analysis on Agilent Mass Profiler Professional software (MPP).

**Targeted metabolomics**

*TCA cycle*

Concentration of TCA analytes were measured by gas chromatograph mass spectrometry (GC-MS) as previously described with a few modifications.^43, 56^ Briefly, 20ul homogenate was lysed by sonication then extracted in 300 ul of chilled methanol and acetonitrile solution after the addition of 10ul of internal solution containing U-^13^C labeled analytes. After drying the supernatant in the speed vac, the sample was derivatized with ethoxime and then with MtBSTFA + 1% tBDMCS (N-Methyl-N-(t-Butyldimethylsilyl)-Trifluoroacetamide + 1% t-Butyldimethylchlorosilane) before it was analyzed on an Agilent 5977B GC/MS (gas chromatography/mass spectrometry) under electron impact and single ion monitoring conditions. Concentrations of lactic acid (m/z 261.2), fumaric acid (m/z 287.1), succinic acid (m/z 289.1), ketoglutaric acid (m/z 360.2), malic acid (m/z 419.3), aspartic acid (m/z 418.2), 2-hydroxyglutaratic acid (m/z 433.2), cis aconitic acid (m/z 459.3), citric acid (m/z 591.4), and isocitric acid (m/z 591.4), glutamic acid (m/z 432.4) were measured against a 7-point calibration curves that underwent the same derivatization.

Acetyl-CoA level

Acetyl CoA was measured by LCMS as previously described.^59^ Briefly, 50uL of homogenates was lysed after the addition of an isotopically labeled acetyl CoA internal standard solution. Protein was removed by adding cold methanol to mixture. The supernatant was transferred to a new vial and dry down prior to resuspending in running mobile phase A for analysis. A 12-point calibration standard curve was constructed using authentic standard and the same internal standard solution as samples. Both standards and samples were analyzed on a Sciex 7500 triple quadrupole mass spectrometer coupled with a Nexera 40 liquid chromatography system. Data acquisition was done using select ion monitor (SRM), m/z 810>303, and 812>305 for acetyl CoA and ^13^C_2_-acetyl CoA respectively via positive electrospray condition.

Glycolysis and pentose phosphate pathway

Tissue was homogenized in 1xPBS after adding 10 µl of PBS to 1mg of tissue prior to aliquoting 50ul homogenate for analysis. Concentration of 10 PPP intermediates were measured against their respective standard curves on GC-MS as previously described.^57^ Briefly, 5 mg of pulverized tissue was homogenized in 1X PBS, spiked with internal standard, and sonicated prior to protein precipitation with cold methanol and acetonitrile mixture (50:50, v:v). The supernatant was dried down and the sample was derivatized to its methoxime-trimethylsilyl derivatives prior to analysis on an Agilent 5977B GC/MS (gas chromatography/mass spectrometry) under electron impact and single ion monitoring conditions. Concentrations of each analyte was measured against its respective 10-point calibration curve that underwent the same derivatization process.

*Acylcarnitine profile*

Tissue was homogenized in 1xPBS after adding 10 µl of PBS to 1mg of tissue prior to aliquoting 25ul homogenate for analysis. Acyl carnitines (specifically C0-C18:1) were measured by liquid chromatography mass spectrometry (LC-MS).^56^ Briefly, homogenate was spiked with a purchased internal standard consisting of isotopically labeled acyl carnitines. The samples were then extracted with cold MeOH:DCM (1:1) followed by centrifugation at 12,000 g for 10 minutes. The supernatant was transferred to another vial, dried down and reconstituted in running buffer. A calibration curve was made from a purchased acyl carnitine mix aliquoted at various concentrations and spiked with the same internal standard as the samples. The samples and calibration standards were analyzed on a triple quadrupole mass spectrometer coupled with an Ultra Pressure Liquid Chromatography system. Data acquisition was done using selective ion monitoring (SRM). The analyte concentrations of each unknown were calculated against their perspective standard curves.

*Amino acid profile*

Amino acids and their metabolites were measured by LC-MS as previously described.^58, 59^ Briefly, 20 µl of homogenate was spiked with an internal standard solution consisting of isotopically labeled amino acids. The supernatant was immediately derivatized with 6-aminoquinolyl-N-hydroxysuccinimidyl carbamate according to Waters’ AccQ-Fluor kit. A 10-point calibration standard curve underwent similar derivatization procedure after the addition of internal standards. Both derivatized standards and samples were analyzed on a Thermo Quantiva triple quadrupole mass spectrometer coupled with a Waters Acquity liquid chromatography system. Data acquisition was done using select ion monitor (SRM) via positive electrospray condition. Concentrations of 42 analytes of each unknown were calculated against its respective calibration curve.
